# Cardiac neurons expressing a glucagon-like receptor mediate cardiac arrhythmia induced by high-fat diet in *Drosophila*

**DOI:** 10.1101/2023.12.13.571403

**Authors:** Yunpo Zhao, Jianli Duan, Joyce van de Leemput, Zhe Han

## Abstract

Cardiac arrhythmia leads to increased risks for stroke, heart failure, and cardiac arrest. Arrhythmic pathology is often rooted in the cardiac conduction system, but the mechanism is complex and not fully understood. For example, how metabolic diseases, like obesity and diabetes, increase the risk for cardiac arrhythmia. Glucagon regulates glucose production, mobilizes lipids from the fat body, and affects cardiac rate and rhythm, attributes of a likely key player. *Drosophila* is an established model to study metabolic diseases and cardiac arrhythmias. Since glucagon signaling is highly conserved, we used high-fat diet (HFD)-fed flies to study its effect on heart function. HFD led to increased heartbeat and an irregular rhythm. The HFD-fed flies showed increased levels of adipokinetic hormone (Akh), the functional equivalent to human glucagon. Both genetic reduction of Akh and eliminating the Akh producing cells (APC) rescued HFD-induced arrhythmia, whereas heart rhythm was normal in Akh receptor mutants (*AkhR^null^*). Furthermore, we discovered a pair of cardiac neurons that express high levels of Akh receptor. These are located near the posterior heart, make synaptic connections at the heart muscle, and regulate heart rhythm. Altogether, this Akh signaling pathway provides new understanding of the regulatory mechanisms between metabolic disease and cardiac arrhythmia.

**HIGHLIGHTS:**

- High-fat diet activates Akh (glucagon-like)-producing neurons near the esophagus in *Drosophila*
- Reducing Akh prevents high-fat diet-induced cardiac arrhythmia in flies
- Discovery of two neurons located at the posterior end of the heart that express the Akh receptor (AkhR) and innervate the heart
- Eliminating one of the two AkhR-expressing cardiac neurons (ACN) results in cardiac arrhythmia, whereas the absence of functional AkhR prevents high-fat diet-induced cardiac arrhythmia in flies

## INTRODUCTION

Arrhythmia refers to an irregular, decreased (bradycardia), or increased (tachycardia) heartbeat. Temporary disruption is usually benign, however chronic arrhythmia has been linked to significantly increased risks for stroke, heart failure, and cardiac arrest (Kannel et al., 1998, 1983; Nattel et al., 2014; Roberts-Thomson et al., 2011). Its pathogenesis is rooted in the cardiac conduction system; however, the mechanism is complex and much remains unknown. Two well-established risk factors that directly contribute to the development of cardiovascular disorders and arrhythmia are obesity (Gupta et al., 2022; Powell-Wiley et al., 2021) and diabetes mellites (Aune et al., 2018; Huxley et al., 2011; Lee et al., 2017). In fact, a longitudinal study into obesity (13.7 years mean follow-up; 5,282 participants) found a 4% increased risk for atrial fibrillation (a form of arrhythmia) per 1-unit increased body mass index (Wang et al., 2004). The other major risk factor, diabetes mellitus has been shown to impact the cardiac conduction system, leading to increased risk of developing atrial fibrillation and ventricular arrhythmias (Kannel et al., 1998). A meta-analysis found that patients with diabetes had a 28-40% increased risk for developing atrial fibrillation, with a 20% risk increase reported for pre-diabetic patients (Aune et al., 2018; Huxley et al., 2011). The relationship between higher blood glucose levels and increased risk for atrial fibrillation was dose-responsive (Aune et al., 2018). Diabetes mellites, when considered a cause of disrupted metabolism, as well as obesity have been independently associated with increased risk for atrial fibrillation (Lee et al., 2017). However, to what extent diabetes, blood glucose, and obesity contribute to atrial fibrillation, independently or collectively, and through which pathomechanism requires further study.

Antagonistic actions by glucagon and insulin regulate glucose metabolism. Besides regulating glucose release from the liver, glucagon facilitates the release of glucose as well as lipids from the fat body, acts as a satiety factor in the central nervous system, affects the glomerular filtration rate, and regulates intra-islet secretion of insulin, glucagon and somatostatin to meet increased energy demands (Habegger et al., 2010; Heppner et al., 2010; Vuguin and Charron, 2011). These glucagon regulatory effects are evident in patient studies that showed that an impaired counter-regulatory glucagon response, observed as increased free plasma insulin levels, contributes to glucose instability in patients with long-term diabetes (Scott et al., 1980); that consumption of dietary fats leads to increased plasma glucagon levels in healthy volunteers (Radulescu et al., 2010); and that plasma glucagon levels were significantly higher in people considered obese compared to those considered lean (Stern et al., 2019). Glucagon has been repeatedly shown to affect heart contraction and heart rate. However, the nature of this effect is complex; whether glucagon acts anti- or pro-arrhythmogenic seems to depend on context, such as a non-failing heart or a heart at acute or chronic failure (Neumann et al., 2023). That said, glucagon-producing tumors, *i.e.*, glucagonomas, can cause tachycardia (a form of arrhythmia defined as >100 heart beats per minute) and heart failure without secondary cause (Chang-Chretien et al., 2004; Zhang et al., 2014). Moreover, glucagonoma tumor resection resulted in a normalized heart rate and a return to typical heart size and function (Chang-Chretien et al., 2004). Similarly, glucagon infusion in healthy human volunteers induced arrhythmias (Jaca et al., 2002; Markiewicz et al., 1978). Thus, while glucagon clearly affects heart rhythm, when considering the underlying mechanism much remains unknown.

The link between high-fat diet (HFD), glucagon, and cardiac arrhythmia is conserved and well-established in animal models. In fact, studies from the 1960s already showed that glucagon increases heart rate in a variety of mammalian species, including dogs, cats, guinea pigs, and rats (Farah and Tuttle, 1960; Lucchesi et al., 1968). More recently, it was shown in mice that both glucagon and the glucagon receptor (Gcgr) are involved in heart rate regulation (Mukharji et al., 2013; Sowden et al., 2007). *Drosophila* adipokinetic hormone (Akh), the functional equivalent to human glucagon, is expressed in a small cluster of endocrine neurons, Akh producing cells (APC) (Kim and Rulifson, 2004; Lee and Park, 2004). These cells in the corpora cardiaca near the esophagus function similar to islet cells in mammals, including the mechanism that regulates hormone secretion, and are essential for larval glucose homeostasis (Kim and Rulifson, 2004). Like glucagon, Akh is known to also mobilize lipids from the fat body to regulate glucose levels, in the form of trehalose, in the circulating hemolymph to accommodate increased energy demand (Isabel et al., 2005). Likewise, increased Akh increases the heart rate in flies (Noyes et al., 1995). These studies together suggest that glucagon and glucagon signaling, aside from regulating blood sugar levels, play an evolutionarily conserved role in heart rate regulation.

*Drosophila melanogaster* is a well-established model to study heartbeat and heart arrhythmia (Birse et al., 2010; Ding et al., 2022; Ocorr et al., 2007). For example, HFD-fed flies were used to study the genetic mechanisms of cardiac dysfunction in obesity. It found that HFD leads to reduced cardiac contractility and a reduced heart period (Birse et al., 2010). Notably, this phenotype was attenuated by intervention at the insulin-TOR signaling pathway (Birse et al., 2010), thus supporting a connection between obesity, glucagon, insulin and cardiac function. Here, we used *Drosophila* fed a HFD to study heart arrhythmia. HFD led to increased Akh in the APC of the corpora cardiaca. We then identified a hitherto undescribed pair of cardiac neurons near the posterior heart that highly express the Akh receptor (AkhR) and directly innervate the fly heart. We show that these AkhR cardiac neurons (ACN) regulate heart rhythm in the flies and mediate the HFD-induced arrhythmia. Given the conservation of Akh/glucagon signaling, these findings likely have implications for arrhythmia in patients. Given the significance of the vagus nerve in cardiac rhythm (Ambache and Lippold, 1949; Cai et al., 2023; Freeman, 1951; González et al., 2023; Kharbanda et al., 2023; van Weperen and Vaseghi, 2023) it could fulfill a similar function.

## RESULTS

### High-fat diet increases heart rate and arrhythmia index

All flies (*w*^1118^) that eclosed (*i.e.*, adults that emerged from pupa) within 8 hours were sorted (females only) and transferred to fresh food vials (25 flies per vial) containing normal fat diet (NFD) or high-fat diet (HFD, NutriFly diet supplemented with 14% fat) for 7 days. At which point, we determined the heartbeat of the flies using optical coherence tomography (OCT) (Figure 1A). Flies on a HFD show a significantly reduced heart period and increased arrhythmia index compared to NFD-fed flies (Figure 1B-D). These findings are in agreement with previous work (Birse et al., 2010) and indicate that HFD imposes a pathogenic effect on heart function.

**Figure 1.**
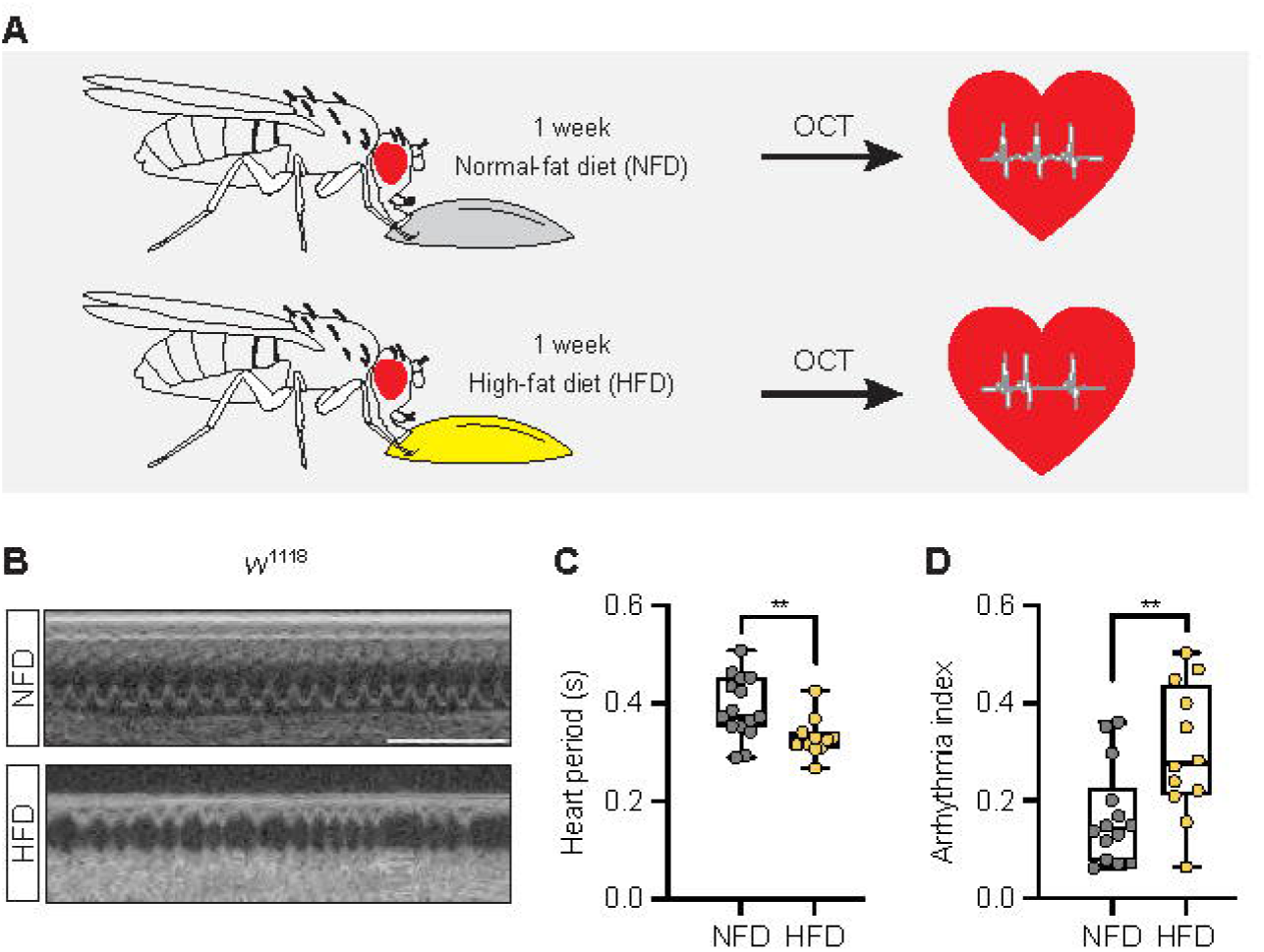
HFD increases heart rate and arrhythmia index in *Drosophila*. **(A)** Schematic illustration of the experimental design. Adult flies (female) were fed either a normal fat diet (NFD) or a high-fat diet (HFD) for 7 days following eclosion from pupa, then subjected to heart functional analysis using optical coherence tomography (OCT) to determine the heart period and the arrhythmia index. **(B)** Representative images obtained from OCT videos of *w*^1118^ flies that were fed NFD or HFD as indicated. Scale bars: 2 seconds. **(C,D)** Quantitation of heart period (**C**; n=15 NFD, n=11 HFD) and arrhythmia index (**D**; n=14 NFD, n=12 HFD). Statistical analysis was performed using *t*-test corrected with Welch; **, P<0.01.

### High-fat diet up-regulates Akh expression and increases Akh producing cell (APC) activity

The ingestion of superfluous macronutrients has a great impact on metabolism and behavior (Huang et al., 2020; Liao et al., 2020; Zhao et al., 2023, 2022). We observed enlarged crops, the functional equivalent to the human stomach, in the HFD-fed flies (Figure 2-figure supplement 1). We asked whether the metabolic signaling pathways contributed to the HFD-caused pathogenic effect on the heart. Akh, the functional equivalent to human glucagon, is expressed by a group of neuroendocrine cells known as APC in the corpora cardiaca (Lee and Park, 2004). Indeed, flies that carried *Akh-*Gal4 to express GFP showed Akh producing cells at the corpora cardiaca, which attaches to the esophagus, just anterior to the proventriculus (Figure 2A). In consistent with a previous work (Liao et al., 2020), we showed that the expression of *Akh* was significantly up-regulated in the flies fed a HFD, compared to NFD-fed flies (Figure 2B).

**Figure 2.**
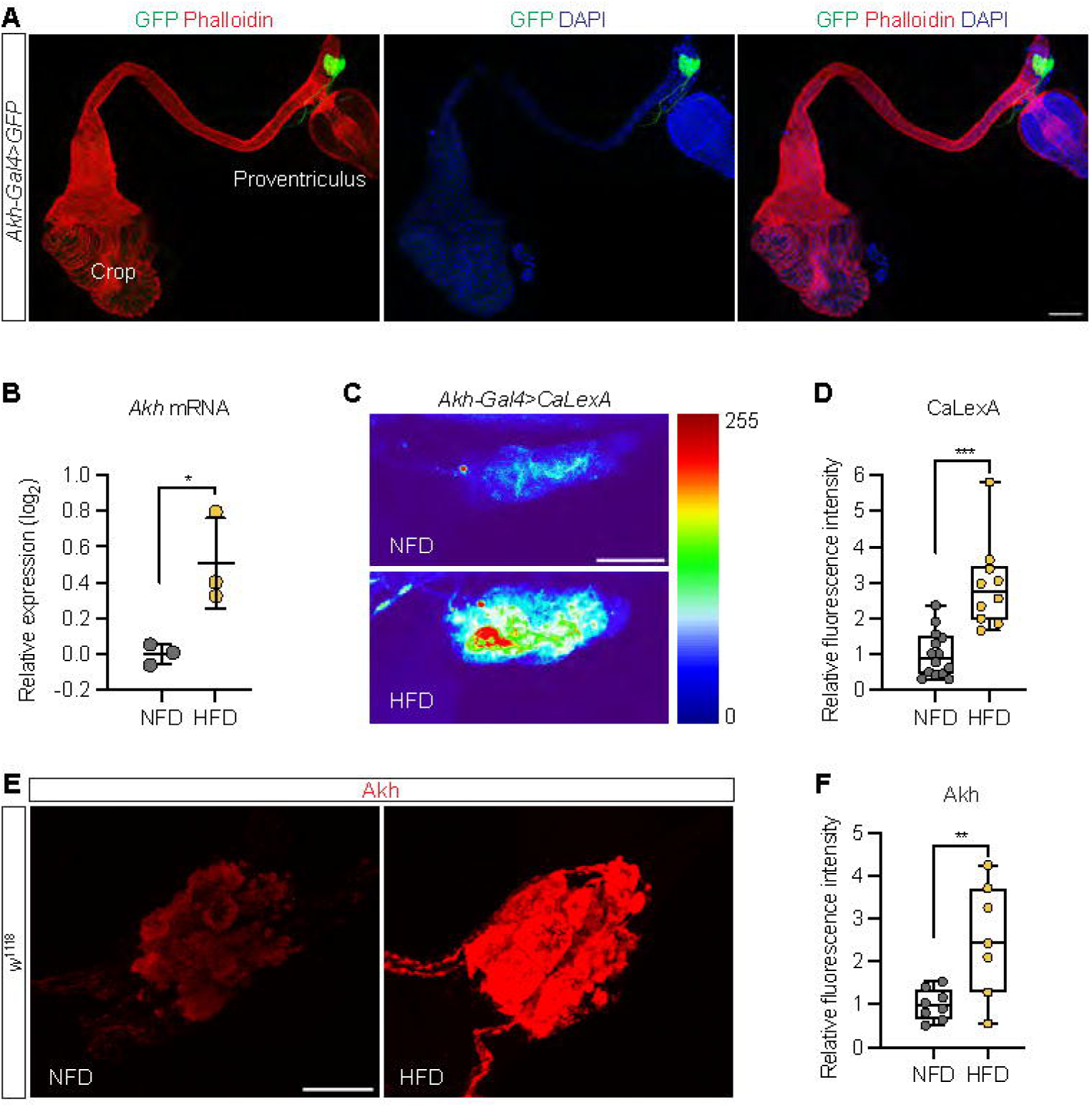
HFD up-regulates adipokinetic hormone gene *Akh* and activates Akh neurons. **(A)** Representative confocal image of Akh producing cells (APCs; green, GFP) in *Akh-*Gal4*>*10xUAS*-GFP* (*Akh*-Gal4>*GFP*) transgenic flies (3-day-old females). The image shows the anterior section of the digestive system, including the crop, esophagus, and proventriculus. The APCs are located at the corpora cardiaca, which attaches to the esophagus, just anterior to the proventriculus. Phalloidin stains actin filaments red; DAPI stains DNA in nucleus blue. Scale bar: 100 μm. **(B)** RT-qPCR analysis of *Akh* expression in *w*^1118^ flies (female) that were fed either a normal fat diet (NFD) or a high-fat diet (HFD) for 7 days following eclosion from pupa. Ten flies per group, repeated three times. Statistical analysis was performed with unpaired *t*-test corrected with Welch; *, P<0.05. **(C)** Representative confocal images of *Akh-*Gal4*>CaLexA* APCs from female adult flies fed an NFD or HFD (for 7 days following eclosion from pupa). CaLexA is a transcription-based genetically encoded calcium indicator for neuronal activity. CaLexA fluorescence has been presented as heatmap; scale from 0 (blue), no CaLexA detected, to 255 (red) high levels CaLexA. Scale bar: 20 μm. **(D)** Quantitation of CaLexA fluorescence in C (7-day-old females; NFD, n=14; HFD, n=10). Statistical analysis was performed with unpaired *t*-test corrected with Welch; ***, P<0.001. (**E**) Representative confocal images of *w*^1118^ APCs from female adult flies fed an NFD or HFD (for 7 days following eclosion from pupa). Anti-Akh is in red. DAPI stains DNA in blue. Scale bar: 20 μm. (**F**) Quantitation of anti-Akh fluorescence in E (7-day-old females; NFD, n=8; HFD, n=7). Statistical analysis was performed with unpaired t-test corrected with Welch; **, P<0.01.

Next, to test whether HFD affects APC neuron activity, we used CaLexA (Masuyama et al., 2012). A basal level of CaLexA fluorescence was observed in the APC of NFD-fed flies, while a significantly higher fluorescence was detected in the APC of HFD-fed flies (Figure 2C, D). This shows that HFD increases APC neuron activity. Taken together, our data show that HFD activates Akh expression in the APC and increases APC activity.

### Akh regulates heartbeat

To confirm the importance of APC activity and Akh release by the APC, we down-regulated *Akh* (*Akh-*Gal4*>*UAS*-Akh*-RNAi) and analyzed its effect on heart function. Immunostaining showed diminished anti-Akh (Lee and Park, 2004) fluorescence (Figure 3-figure supplement 1), indicating the RNAi efficiency. As expected, upon Akh depletion in the APCs, the difference in arrhythmia between the NFD and HFD-fed flies disappeared (Figure 3A, B). The findings indicate that the endocrine signal Akh, originating in the APC neurons, mediates the HFD cardiac functional pathogenicity.

**Figure 3.**
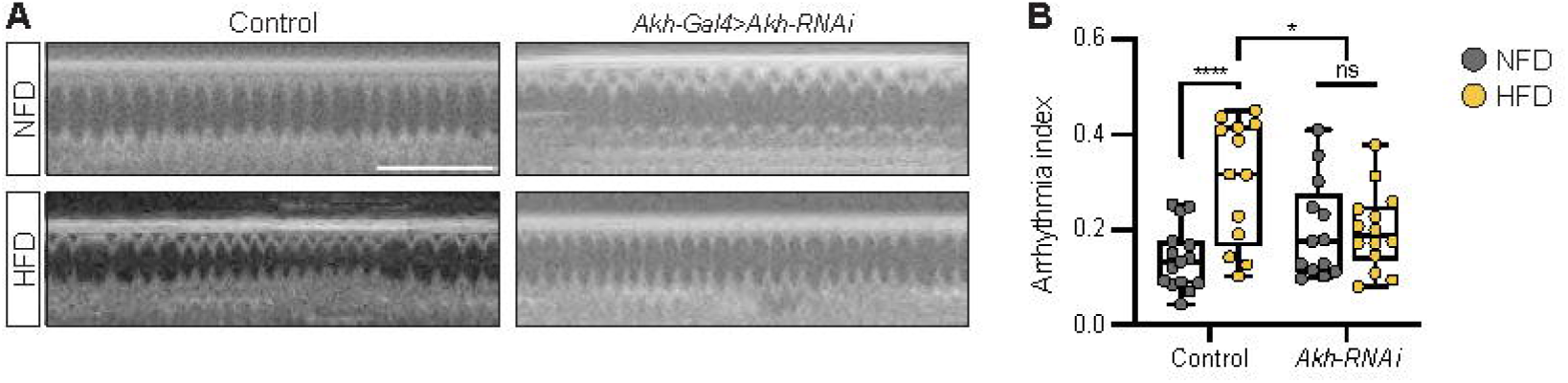
Akh regulates the heartbeat. **(A)** Representative images obtained from optical coherence tomography (OCT) videos of control (*Akh*-Gal4) and *Akh*-RNAi (*Akh*-Gal4>UAS-*Akh*-RNAi) flies (females, 7 days old) that were fed a normal fat diet (NFD) or a high-fat diet (HFD), as indicated, for seven days starting at eclosion from pupa. Scale bars: 2 seconds. **(B)** Quantitation of the arrhythmia index in control (n=15 NFD, n=13 HFD) and *Akh*-RNAi (n=13 NFD, n=14 HFD) flies. Statistical analysis was performed with two-way ANOVA corrected with Sidak. Statistical significance: *, p<0.05; ****, p<0.0001; ns, not significant.

### AkhR mediates the high-fat diet pathogenic effect on the heartbeat

Akh binds its receptor AkhR, a G-protein coupled receptor, to activate the signaling pathway (Staubli et al., 2002). We quantified *AkhR* expression using RT-qPCR and observed significantly up-regulated expression in HFD-fed flies (Figure 4A). To confirm its role in Akh-mediated HFD-induced arrhythmia, we tested *AkhR* null mutant flies (*AkhR^null^*). In line with Akh depletion, flies with *AkhR* mutation and fed a HFD, did not show the HFD-associated cardiac arrhythmia (Figure 4B, C). These data show that Akh/AkhR signaling mediates the pathogenic effect of HFD on heart function.

**Figure 4.**
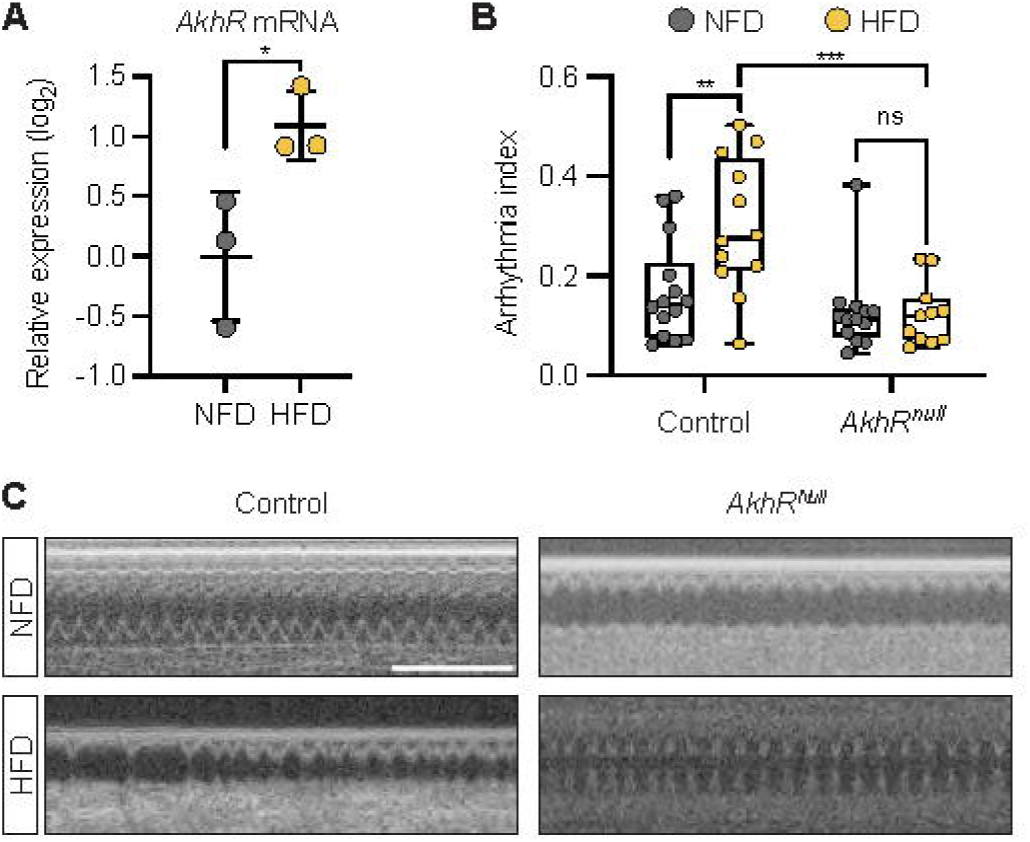
AkhR regulates the heartbeat. **(A)** RT-qPCR analysis of *AkhR* expression in *w*^1118^ flies (female) that were fed either a normal fat diet (NFD) or a high-fat diet (HFD) for 7 days following eclosion from pupa. Ten flies per group, repeated three times. Statistical analysis was performed with unpaired *t*-test corrected with Welch; *, P<0.05. **(B)** Quantitation of arrhythmia index in control (n=14 NFD, n=12 HFD) and *AkhR^null^* (n=11 NFD, n=13 HFD) flies. Statistical analyses were performed with two-way ANOVA corrected with Sidak; **, P<0.01; ***, P<0.001; ns, not significant. **(C)** Representative images obtained from optical coherence tomography (OCT) videos of control (*w*^1118^) and *AkhR^null^* mutant flies flies (females, 7 days old) that were fed NFD or HFD, as indicated, for seven days starting at eclosion from pupa. Scale bars: 2 seconds.

### A pair of AkhR cardiac neurons (ACN) are associated with the heart

To determine the AkhR expression pattern, we used *AkhR-*Gal4 (Lee et al., 2018) to drive the expression of GFP. The adipose fat body tissue is a major organ that expresses AkhR at the embryonic and larval stages (Grönke et al., 2007), as well as in the adult (Bharucha et al., 2008). Likewise, we observed GFP fluorescence in the fat body in *AkhR-*Gal4*>GFP* flies (Figure 5A, B). Notably, no fluorescence was detected in the cardiac muscles. However, we did find two neurons with strong GFP fluorescence, indicative of high expression levels of AkhR, located near the posterior end of the heart tube (Figure 5B). These neurons had elaborate neurites along the heart tube (Figure 5B-D) and formed synaptic connections with heart muscles, as revealed by immunostaining for active zone marker Bruchpilot (Brp) (Figure 5E, F). These two neurons likely communicate with the heart muscle via these neuromuscular junctions. Therefore, were refer to these two neurons as AkhR cardiac neurons (ACN). Immunostaining for Akh showed positive fluorescence on the ACN (Figure 5-figure supplement 1), suggesting that the cardiac neurons receive Akh signal.

**Figure 5.**
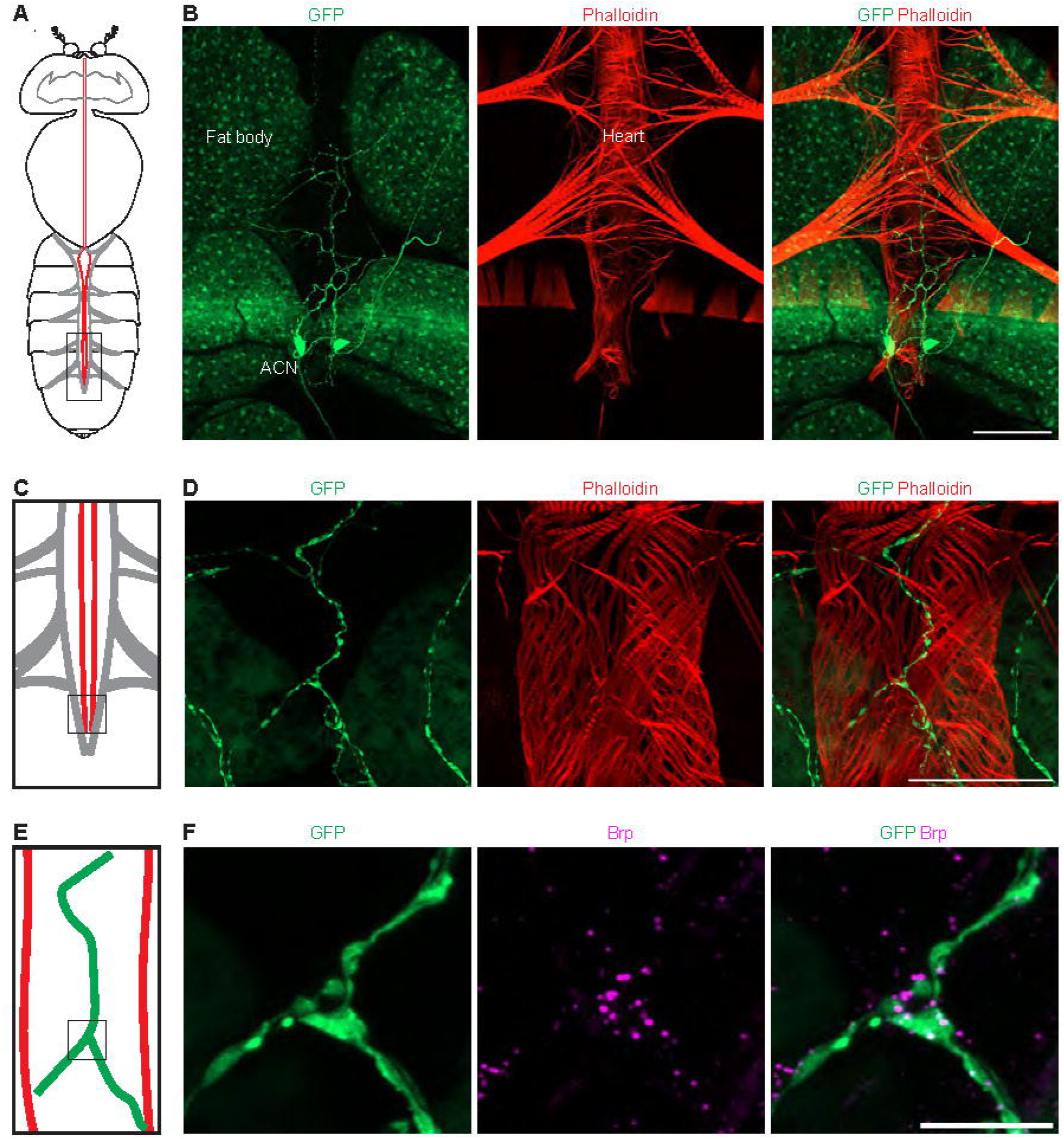
AkhR cardiac neurons (ACNs) form synaptic connections with the posterior heart. **(A)** Graphic depiction of the body (no legs or wings) of an adult fly with the head oriented towards the top. The heart tube (red) is located along the midline in the abdomen (bottom of graphic), with the alary muscles that connect the heart to the exoskeleton represented in light grey. **(B)** A representative confocal image of an *AkhR-*Gal4*>*10xUAS*-GFP* adult fly (7-day-old, female). The dorsal region of an abdomen corresponding to the boxed region in (A) is shown. *AkhR*>*GFP* labels AkhR in green. ACN, AkhR cardiac neuron. Phalloidin stains actin filaments red. Scale bar: 100 μm. **(C)** Schematic illustration of the posterior fly heart. Corresponds to boxed region in (A). **(D)** A representative confocal image of the posterior heart of an *AkhR-*Gal4*>*10xUAS*-GFP* fly (7-day-old, female) corresponding to the boxed region in (C). *AkhR*>*GFP* labels AkhR in green. Phalloidin stains actin filaments red. Scale bar: 50 μm. **(E)** Schematic illustration of the posterior end of an adult fly heart. The neurites of the cardiac neurons are depicted in green. Relates to the boxed region in (C). **(F)** Confocal image of the posterior heart of an *AkhR-*Gal4*>*10xUAS*-GFP* fly (7-day-old, female) corresponding to the boxed region in (E). *AkhR*>*GFP* labels AkhR in green. Anti-Bruchpilot (Brp), a marker for the active zone, is shown in magenta. Scale bar: 10 μm.

### Partial elimination of AkhR cardiac neurons (ACN) causes arrhythmia

To determine the function of the ACN, we set out to eliminate the pair. We overexpressed UAS-*rpr* under control of *AkhR*-Gal4 to induce apoptosis. We observed one remaining AkhR cardiac neuron in the *AkhR*-Gal4>UAS-*rpr* flies (Figure 6A), indicating partial elimination. The *AkhR*-Gal4>UAS-*rpr* flies were subjected to OCT analysis. The profile and rhythm of the heartbeat were drastically affected in the flies with only one ACN (Figure 6B). This demonstrates the importance of the ACN in regulating the heartbeat.

**Figure 6.**
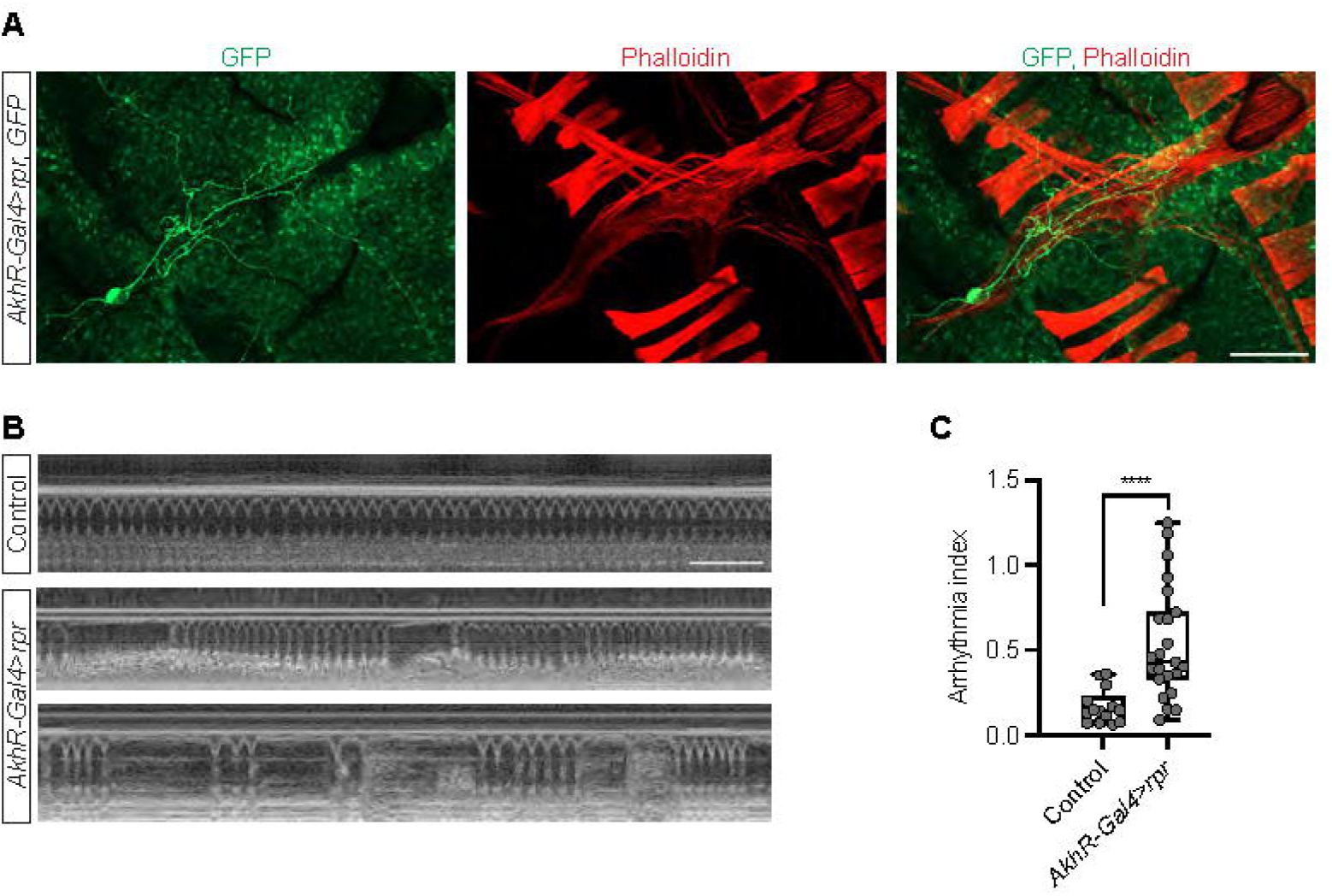
Partial elimination of ACNs causes arrhythmia. **(A)** A representative confocal image of an adult *AkhR*-Gal4>*rpr*, *GFP* (*AkhR-*Gal4*>*UAS*-rpr*, UAS*-GFP*) fly (7-day-old, female). *AkhR*>*GFP* labels AkhR in green. *AkhR*, *adipokinetic hormone receptor*; *rpr*, *reaper*. Phalloidin stains actin filaments red. Scale bar: 100 μm. **(B)** Representative images obtained from optical coherence tomography (OCT) videos of control (*w*^1118^) and *AkhR*-Gal4>*rpr* (*AkhR-*Gal4*>*UAS*-rpr*, UAS*-GFP*) flies (7-day-old, female). Scale bar: 2 seconds. **(B)** Quantitation of arrhythmia index in control (n=14 NFD) and *AkhR*-Gal4>*rpr* (n=23 NFD). Statistical analysis was performed using *t*-test corrected with Welch; ****, P<0.0001.

## DISCUSSION

### The role of AkhR, glucagon-like receptor, in regulating heart rate and rhythm

Heart function is dependent on ATP continuous synthesis. Cardiac ATP comes from fatty acids, glucose, and lactate (Kodde et al., 2007). Glucagon converts stored glycogen, in the liver and fat tissue, into glucose that is released into the blood stream/ Its link to cardiac arrhythmia has been well-established. The glucose signaling pathway is conserved across species, including *Drosophila* (Kim and Rulifson, 2004). Like in humans, in flies the glucagon-like hormone Akh regulates the glucose levels in hemolymph (fly equivalent to blood) (Kim and Rulifson, 2004). Under starvation conditions, Akh mobilizes glycogen and lipids to maintain the circulatory glucose levels. The flies become hyperactive and show increased food searching behavior. Flies without Akh, either by depletion or APC elimination, show starvation resistant behavior, suggesting that the behavioral change is mediated by Akh (Huang et al., 2020; Lee and Park, 2004; Yu et al., 2016). Notably, in flies fed a HFD, the AkhR expressing neurons in the brain become hypersensitive to Akh stimulation, due to upregulated AkhR (Huang et al., 2020). Finally, prepupae injected with low pharmacological doses of Akh, *i.e.*, doses higher than physiologically expected, showed increased heart rates (Noyes et al., 1995). Thus, in both humans and flies, heart rate and rhythm respond to nutritional changes (flies, see Figure 1), this mechanism likely supports the higher metabolic demands and possibly mediates the switch from glucose to stored lipids as the main energy source.

To date, studies have focused on the direct effect of glucagon on cardiac tissue, based on the notion that cardiomyocytes express glucagon receptors (GCGR). However, evidence of GCGR expression in human heart tissue has been conflicting. No cardiac signal was detected when using radioactively labeled glucagon (Bomholt et al., 2022), results of mRNA data have been inconsistent (Aranda-Domene et al., 2023; Bomholt et al., 2022), and protein data for GCGR to support cardiac expression is lacking (Neumann et al., 2023). Here, we identified an endocrine neuron-heart axis using a fly model for high-fat diet-induced arrhythmia. In this fly model (Figure 7), the consumption of a HFD elevates the expression of Akh, a glucagon-like hormone, and activity of the APC. The elevated circulatory Akh is transported to the AkhR cardiac neurons (ACN) located near the posterior end of the heart, where the increased signaling leads to an escalated heart arrhythmia index. We speculate that the ACNs are sympathetic in the adults, which is in line with Akh functioning as a cardioaccelerator in prepupae (Noyes et al., 1995). These findings revealed a hitherto undescribed regulatory signaling pathway that links the consumption of superfluous macronutrients with heart arrhythmia.

**Figure 7.**
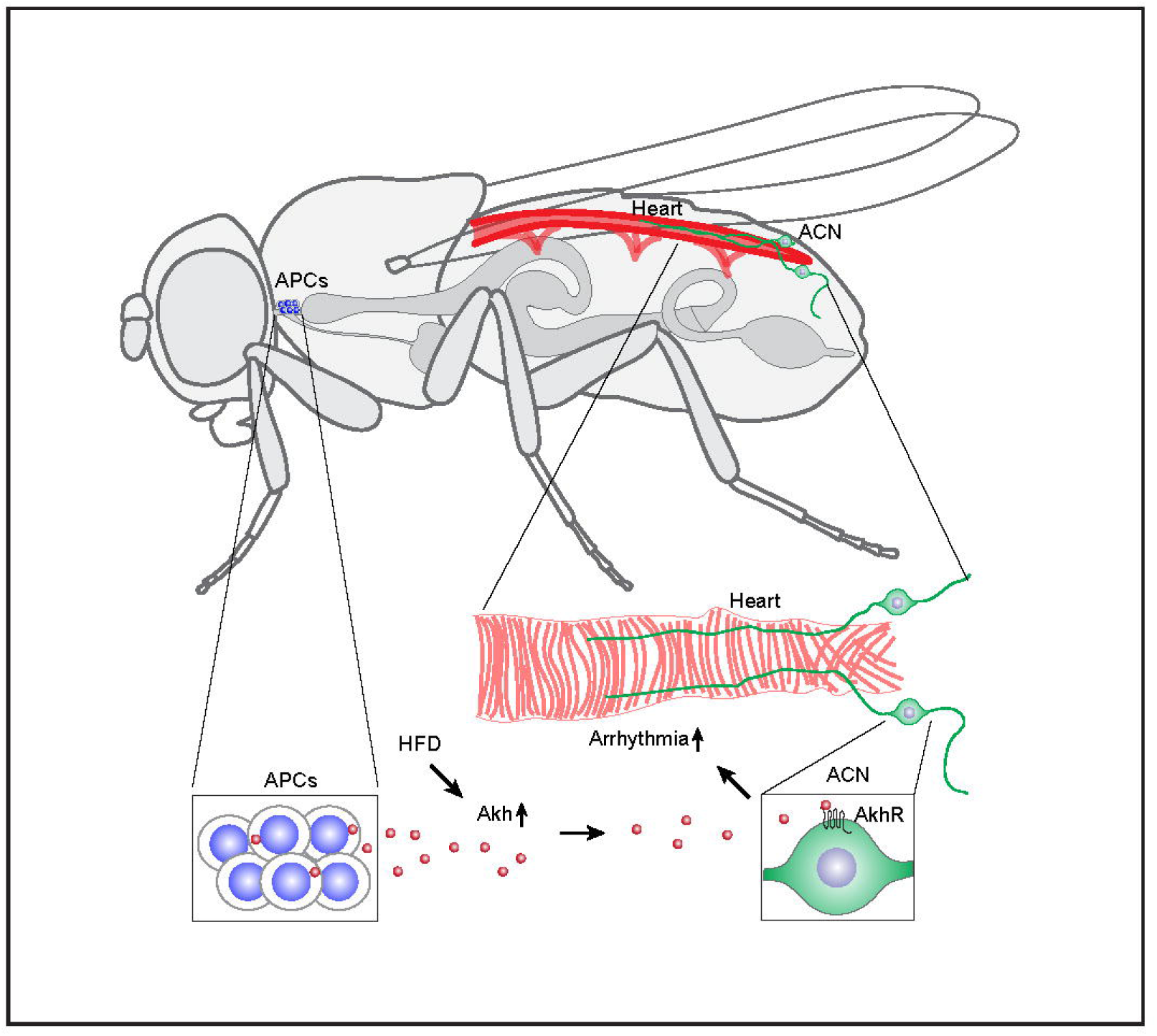
Model: Akh/AkhR mediates the HFD-induced cardiac arrhythmia. A high-fat diet (HFD) upregulates adipokinetic hormone (Akh) levels in the Akh producing cells (APCs) located at the corpora cardiaca adjacent to the anterior esophagus. HFD also increases APC activity, this results in increased Akh secretion into the circulation. Akh acts on the Akh receptors (AkhR) expressed by the AkhR cardiac neurons (ACN). The ACN are a pair of neurons located at the posterior end of the adult fly heart, that regulate the heartbeat. Through this signaling pathways, increased Akh under HFD conditions leads to increased arrhythmia.

### Cardiac regulatory neurons in *Drosophila*

*Drosophila* have a cardiac cycle that consists of alternating retrograde and anterograde heart beats that correlate to the multi-chamber diastole and systole, respectively (Dulcis and Levine, 2005). Accordingly, two signaling systems that innervate the fly heart were identified. The retrograde beat being regulated by glutamatergic innervation. Its transverse nerves run bilaterally along the longitudinal muscle and innervate the cardiac muscles of the conical chamber as well as the alary muscles (Dulcis and Levine, 2005, 2003). The anterograde heartbeat being regulated by crustacean cardioactive peptide (CCAP) innervation. Its bipolar neurons (BpN) extent CCAP fibers that innervate each segment of the abdominal heart. Four additional large CCAP-positive neurons innervate the terminal chamber (Dulcis et al., 2005; Dulcis and Levine, 2003). These earlier studies, leave the possibility of synaptic input or hormonal regulation of the BpN neurosecretory signals to the heart. The authors comment on the absence of direct descending input from the corpora cardiaca to the *Drosophila* abdominal heart as has been observed in other systems (Dulcis and Levine, 2003). Here, we identified corpora cardiaca released Akh signaling to previously unreported AkhR cardiac neurons (ACN) that could exercise this function in flies. Given their localization at the terminal end of the cardiac chamber, where the four large CCAP-positive neurons are located as well, it is possible that the ACN modulate CCAP signaling at the heart. In fact, the previous study detected synaptotagmin on the BpN (BpN6) cell bodies which suggests the presence of presynaptic sites (Dulcis and Levine, 2003). We are currently investigating if the BpN and ACN act together, whether in concert or opposingly.

### Relevance of findings in fly for humans

This study revealed how glucagon-like hormone Akh is released by APCs in response to HFD and stimulates AkhR cardiac neurons (ACN) to regulate heart rhythm in flies. Like in the flies, glucagon infusion in healthy human volunteers induces arrhythmias (Jaca et al., 2002; Markiewicz et al., 1978). Moreover, a preliminary study in infants and children demonstrated the potential of glucagon to treat atrioventricular (AV) block, a heart rhythm disorder marked by a slow heart rate caused by dysfunctional electrical conduction (Hurwitz, 1973). Clinical guidelines by the American Heart Association recommend the use of glucagon to treat bradycardia due to beta-blocker or calcium channel blocker overdose (Kusumoto et al., 2019). However, the mechanism by which glucagon exerts these beneficial clinical effects remains poorly understood. Our findings implicate a potentially conserved signaling pathway in which elevated glucagon leads to increased cardiac neuronal activity and subsequent increased heart rate and arrhythmia. Multiple trials are investigating the possible benefits of glucagon receptors (GCGR) agonists alone or in combination with GLP-1 agonists to treat arrhythmia, but so far the results have been mixed. An initial trial with a GCGR antagonist in patients with type 2 diabetes was halted prematurely due to the detrimental side effects, which included elevated blood cholesterol and steatosis (liver dysfunction) (Agrawal and Gupta, 2016; Kazda et al., 2016a, 2016b; Pearson et al., 2016). Our findings provide new possible glucagon-related targets for treating obesity-associated cardiac arrhythmia by targeting cardiac neurons receptive to glucagon signaling rather than the cardiomyocytes as had been the leading hypothesis to date. Studies into the human equivalent of ACN would be interesting; possibly the vagus nerve could fulfill a similar function as its importance for cardiac rhythm has been firmly established (Ambache and Lippold, 1949; Cai et al., 2023; Freeman, 1951; González et al., 2023; Kharbanda et al., 2023; van Weperen and Vaseghi, 2023).

Finally, while the exact implications of these findings for new therapeutics remain to be seen, they do support the avoidance of a high-fat diet. If unavoidable, for example in the case of a therapeutic ketogenic diet or in the subset of patients with diabetes that have elevated glucagon blood levels (D’Alessio, 2011; Unger et al., 1970), then targeting the glucagon pathway might protect against harmful cardiac side effects.

## MATERIALS AND METHODS

### Key resources table

| Reagent type (species) or resource | Designation | Source or reference | Identifiers | Additional information |
| --- | --- | --- | --- | --- |
| Antibody | anti-Brp (mouse IgG1, kappa light chain) | Developmental Studies Hybridoma Bank | Cat# Nc82 RRID: AB_2314866 | IF (1:100) |
| Antibody | goat anti-mouse Alexa Fluor 647 | Jackson Immunoresearch Laboratories | Cat# 115-605-003 | IF (1:500) |
| Antibody | anti-Akh (rabbit) | Prof. Jae Park | N.A. | IF (1:1000) |
| Antibody | goat anti-rabbit Alexa Fluor 568 | ThermoFisher | Cat# A-11011 | IF (1:500) |
| Other | Alexa Fluor 647 Phalloidin | ThermoFisher | Cat# A22287 | IF (1:1000) |
| Other | normal goat serum | Jackson Immunoresearch Laboratories | Cat# 102643-594 | 1% in 1xPBST |
| Genetic reagent (*Drosophila melanogaster*) | *w*<sup>1118</sup> | Bloomington Drosophila Stock Center | BDSC_3605 |  |
| Genetic reagent (*Drosophila melanogaster*) | *Akh*-Gal4 | Bloomington Drosophila Stock Center | BDSC_25684 |  |
| Genetic reagent (*Drosophila melanogaster*) | UAS-*Akh*-RNAi | Bloomington Drosophila Stock Center | BDSC_34960 |  |
| Genetic reagent (*Drosophila melanogaster*) | *w*<sup>1118</sup> P{UAS-rpr.C}27; P{UAS-2xEGFP}AH3 | Bloomington Drosophila Stock Center | BDSC_91417 |  |
| Genetic reagent (*Drosophila melanogaster*) | *AkhR*<sup>null</sup> | Bloomington Drosophila Stock Center | BDSC_80937 |  |
| Genetic reagent (*Drosophila melanogaster*) | *w*[*]; P{w[+mC]=LexAop-CD8-GFP-2A-CD8-GFP}2; P{w[+mC]=UAS-*mLexA-VP16-NFAT*}H2, P{w[+mC]=lexAop-rCD2-GFP}3/TM6B, Tb[1] | Bloomington Drosophila Stock Center | BDSC_66542 |  |
| Genetic reagent (*Drosophila melanogaster*) | 10xUAS-*GFP* | Bloomington Drosophila Stock Center | BDSC_32185 |  |
| Genetic reagent (*Drosophila melanogaster*) | *AkhR-T2A*-Gal4 | Bloomington Drosophila Stock Center | BDSC_78877 |  |
| Chemical compound, drug | 4% paraformaldehyde (PFA) | Thermo Fisher Scientific | Cat# J19943.K2 |  |
| Chemical compound, drug | Triton X-100 | Sigma-Aldrich | Cat# 9002-93-1 | 0.2% in 1xPBS |
| Chemical compound, drug | 4',6-Diamidino-2-phenylindole (DAPI) | Thermo Fisher Scientific | Cat# D1306 RRID: AB_2629482 | (0.5 µg/ml) |
| Commercial assay or kit | 10x phosphate buffered saline | Quality Biological | Cat# 119-069-131 |  |
| Commercial assay or kit | TRIzol | Thermo Fisher Scientific | Cat# 15596026 |  |
| Commercial assay or kit | EcoDry Premix kit | Takara Bio | Cat# 639549 |  |
| Commercial assay or kit | PowerSYBR Green | Applied Biosystems - Thermo Fisher Scientific | Cat# 4367659 |  |
| Commercial assay or kit | VECTASHIELD | Vector Laboratories | Cat# H-1000 |  |
| Software, algorithm | Adobe Illustrator 2022 | Adobe Inc. | RRID:SCR_010279 |  |
| Software, algorithm | ImageJ | National Institutes of Health | RRID:SCR_003070 |  |
| Software, algorithm | GraphPad Prism9 | GraphPad | RRID:SCR_002798 |  |

### Drosophila husbandry

*Drosophila* were reared on a normal fat diet (NFD, Nutri-Fly German formula; Genesee Scientific, San Diego, CA; 66-115) or a high-fat diet (HFD, NFD supplemented with 14% coconut oil), under standard conditions (25°C, 60% humid, 12h:12h dark:light). The following fly lines were obtained from Bloomington Drosophila Stock Center (BDSC) at Indiana University Bloomington (Bloomington, IN): *w*^1118^ (BDSC_3605), *Akh-*Gal4 (BDSC_25684), UAS*-Akh-*RNAi (BDSC_34960), *Akh-T2A-*Gal4 (BDSC_78877), *w*^1118^ P{UAS-*rpr.C*}27; P{UAS-2x*EGFP*}AH3 (BDSC_91417), *AkhR^null^* (BDSC_80937), *w[*]; P{w[+mC]=LexAop-CD8-GFP-2A-CD8-GFP}2; P{w[+mC]=*UAS*-mLexA-VP16-NFAT}H2, P{w[+mC]=lexAop-rCD2-GFP}3/TM6B, Tb[1]* (BDSC_66542), and *10x*UAS*-GFP* (BDSC_32185).

### Optical coherence tomography (OCT)

Cardiac function in adult *Drosophila* was measured using an OCT system (Bioptigen; this system was built by the Biophotonics Group, Duke University, Durham, NC (Yelbuz et al., 2002)). For this, newly eclosed adult flies (female) were transferred to fresh vials containing NFD or HFD for 7 days. The flies were anesthetized by carbon dioxide (CO_2_) and mounted onto a glass slide using fixogum rubber cement (Marabu GmbH & Co. KG, Tamm, Germany; MR290117000). The flies were allowed to recover for 10 minutes and then imaged using OCT. The flies were scanned at the cardiac chamber in abdominal segment A2. The acquisition parameters were as follows: 48 Hz, 800 frames.

### Quantification of heart period and arrhythmia index

The heart period and arrhythmia index were quantified as previously reported (Ocorr et al., 2007) with minor modifications. In brief, the OCT raw data was processed using ImageJ software (version 2.9.0/1.53) (Schneider et al., 2012). Within each OCT image (10 seconds) the heart periods were arranged by length. The heart period value for each fly was based on the average of three short, three medium and three long heart periods (*i.e.*, nine periods in total). The arrhythmia index for a genotype was based on the group average (7-day-old females; n=11-23), presented with the standard deviation (SD).

### Immunostaining and confocal imaging

Flies were dissected in 1xPBS, then fixed in 4% PFA for 30 at room temperature. The specimens were then washed in PBST (0.2% Triton X-100 in 1xPBS) three times, 15 min each, followed by blocking in 1% normal goat serum (Jackson ImmunoResearch Laboratories, West Grove, PA; 102643-594) in PBST for 1 hour at room temperature. The specimens were incubated in primary antibody at 4°C overnight, then washed in PBST three times 15 min. Next, the specimens were incubated in secondary antibody for 2 hours at room temperature, followed by three 15 min washes in PBST, followed by a 10 min wash in PBS. DAPI staining (Thermo Fisher Scientific, Waltham, MA; D1306, 0.5 μg/ml in PBST) was performed in-between the washing steps after the secondary antibody staining. The following antibodies were used: mouse monoclonal anti-Brp 1:100 (Developmental Studies Hybridoma Bank, Iowa City, IA; nc82, RRID:AB_2314866); rabbit anti-Akh (1:1000); goat anti-rabbit Alexa Fluor 568 1:500 (ThermoFisher; A-11011); goat anti-mouse Alexa Fluor 647 1:500 (Jackson ImmunoResearch Laboratories, West Grove, PA; 115-605-003). Alexa Fluor 647 Phalloidin 1:1,000 (Thermo Fisher Scientific, Waltham, MA; A22287) was used to stain the filament actin. Antibodies were diluted in the blocking buffer. The specimens were mounted with Vectashield antifade mounting media (Vector Laboratories, Newark, CA; H-1000). The samples were imaged using a ZEISS LSM 900 confocal microscope and ZEISS ZEN blue edition (version 3.0) acquisition software under a 20x Plan-Apochromat 0.8 N.A. air objective or a 63x Plan-Apochromat 1.4 N.A. oil objective. For quantitative comparison of intensities, settings were chosen to avoid oversaturation (using range indicator in ZEN blue) and applied across images for all samples within an assay. Image J was used for image processing (version 2.9.0/1.53t; National Institutes of Health, Bethesda, MD) (Schneider et al., 2012).

### Reverse Transcriptase-quantitative PCR (RT-qPCR)

For quantification of Akh and AkhR mRNA levels, adult female flies fed NFD or HFD for 7 days were transferred into 1.5 mL Eppendorf tubes (10 flies/tube) and flash frozen in liquid nitrogen. Total RNA was extracted using TRIzol (Thermo Fisher Scientific, Waltham, MA, Cat# 15596026) according to the manufacturer’s protocol. In brief, 0.5 mL TRIzol was added to each tube, then flies were grinded using a pestle. Following, 0.1 ml chloroform was added, sample tubes securely capped, and vigorously shaken by hand for 15 seconds. Samples were incubated at 22°C for 2 minutes, then centrifuged at 12,000x*g* for 15 minutes at 4°C, and supernatants transferred to a clean tube in which total RNA was precipitated using isopropanol (Sigma-Aldrich, St. Louis, MO; 67-63-0). Synthesis of cDNA using reverse transcriptase was performed with the RNA to cDNA EcoDry Premix kit (Takara Bio, San Jose, CA; 639549) and and the subsequent quantitative PCR using PowerSYBR Green (Applied Biosystems - Thermo Fisher Scientific, Waltham, MA; 4367659), according to the manufacturers’ protocols. qPCR was performed using a QuantStudio 6 Pro Real-Time PCR machine (Thermo Fisher Scientific, Waltham, MA). The following primers (5’-3’) were used: Akh_F TTTCGAGACACAGCAGGGCA, with Akh_R GGTGCTTGCAGTCCAGAAA; AkhR_F ACAATCCGTCGGTGAACA, with AkhR_R CATCACCTGGCCTCTTCCAT. For each treatment, three biological replicates were prepared. ΔΔCT method was used to quantify the gene expression levels with normalization to *Ribosomal protein L32* (*RpL32*) as internal reference gene (Rpl32_F AACCGCGTTTACTGCGGCGA, with Rpl32_R AGAACGCAGGCGACCGTTGG).

### CaLexA assays

CaLexA assays were performed to determine the neuron activity. *Akh-*Gal4 virgins were crossed with CaLexA males (*w[*]; P{w[+mC]=LexAop-CD8-GFP-2A-CD8-GFP}2; P{w[+mC]=*UAS*-mLexA-VP16-NFAT}H2, P{w[+mC]=lexAop-rCD2-GFP}3/TM6B, Tb[1]*) and maintained on NFD. The newly eclosed adults were transferred to fresh food vials containing NFD or HFD for 3 days. The flies were then dissected in 1x phosphate buffered saline (1xPBS) (Quality Biological, Gaithersburg, MD; 119-069-131) at room temperature and fixed in 4% paraformaldehyde (PFA) (Thermo Fisher Scientific, Waltham, MA; J19943.K2) in 1xPBS for 1 hour at room temperature, followed by three 15 min washes in 1xPBS with Triton X-100 (Sigma-Aldrich, St. Louis, MO; 9002-93-1) (PBST). The specimens were then stained in DAPI (0.5 μg/ml in PBST) (Thermo Fisher Scientific, Waltham, MA; D1306) for 10 min and washed once for 15 min in 1xPBS before mounting with Vectashield antifade mounting media (Vector Laboratories, Newark, CA; H-1000). The samples were imaged using a ZEISS LSM 900 confocal microscope and ZEISS ZEN blue edition (version 3.0) acquisition software under a 20x Plan-Apochromat 0.8 N.A. air objective a 63x Plan-Apochromat 1.4 N.A. oil objective. For quantitative comparison of intensities, settings were chosen to avoid oversaturation (using range indicator in ZEN blue) and applied across images for all samples within an assay. Image J was used for image processing (version 2.9.0/1.53t; National Institutes of Health, Bethesda, MD) (Schneider et al., 2012).

### Data analysis and figure preparation

Figures were arranged using Adobe Illustrator software (version 26.2.1; Adobe Inc., San Jose, CA). The relative fluorescence intensity was acquired using ImageJ software (version 2.9.0/1.53t; National Institutes of Health, Bethesda, MD) (Schneider et al., 2012). Data plotting and statistical analyses were performed using Prism 9 (GraphPad Software, Boston, MA). Data normality was tested by using the Shapiro-Wilk test. Normally distributed data were analyzed by Student’s *t*-test with Welch’s correction (two groups) or by a one-way ANOVA followed by Dunnett’s correction or two-way ANOVA corrected with Sidak. P-value < 0.05 was considered significant.

## ACKNOWLEDGMENTS

We would like to thank the Bloomington Drosophila Stock Center (BDSC) based at Indiana University Bloomington (Bloomington, IN) for the *Drosophila* stocks, Prof. Jae Park (The University of Tennessee) for providing the antibodies, and the Developmental Studies Hybridoma Bank (DSHB) based at the University of Iowa (Iowa City, IA) for providing the antibodies.

## DATA AVAILABILITY

All relevant data can be found within the article and its supplementary information.

## AUTHOR CONTRIBUTIONS

YZ and HZ designed the project; YZ and JD carried out the experiments; YZ carried out the data analyses; YZ prepared the figures; YZ and JvdL wrote the draft and revisions; HZ provided project oversight, resources, and funding. The manuscript has been critically reviewed and the final version approved by all authors.

## FIGURE SUPPLEMENT

**Figure 2—figure supplement 1.**
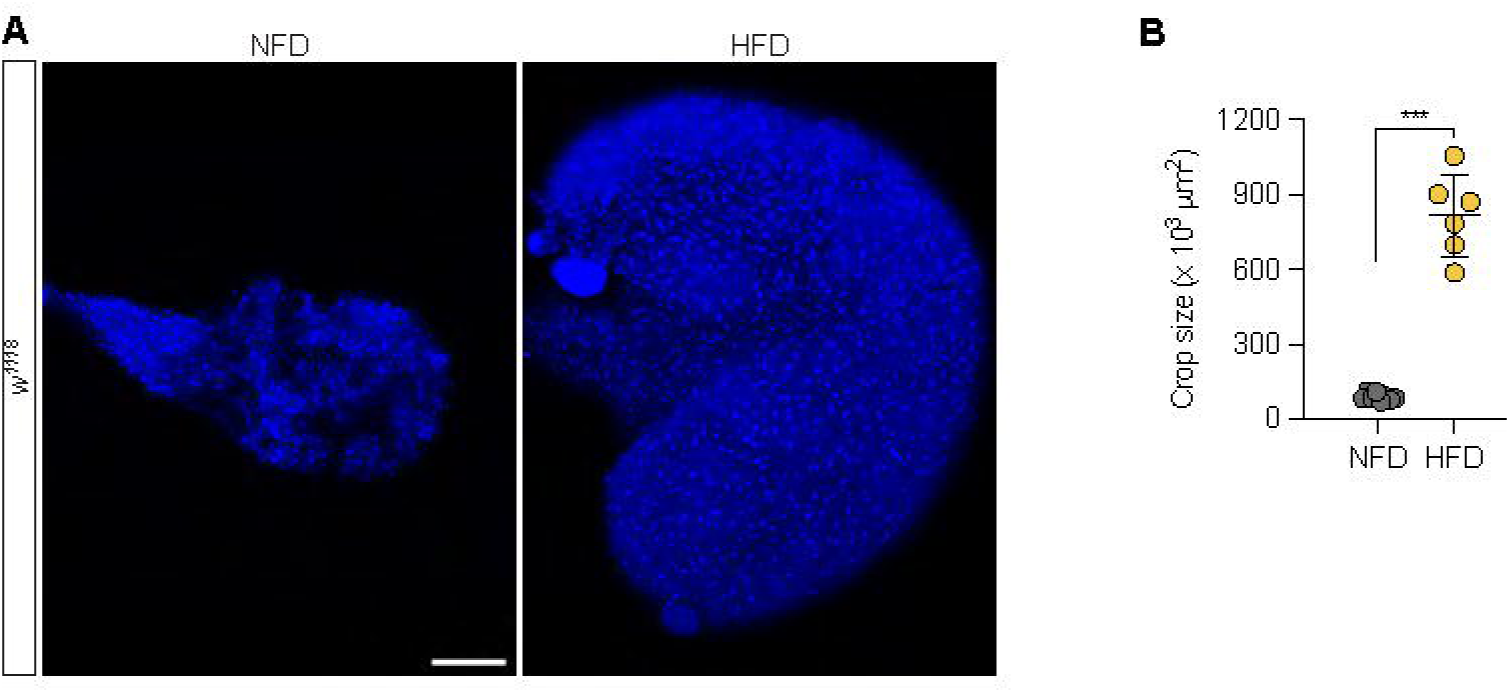
HFD increases crop size. **(A)** Representative confocal images of crops from female adults (*w*^1118^) fed a normal fat diet (NFD) or a high-fat diet (HFD) for 7 days following eclosion from pupa. DAPI stains DNA in blue. Scale bar: 100 μm. **(B)** Quantitation of crop area. n=8 NFD and n=6 HFD. Statistical analysis was performed using *t*-test corrected with Welch; ***, P<0.001.

**Figure 3—figure supplement 1.**
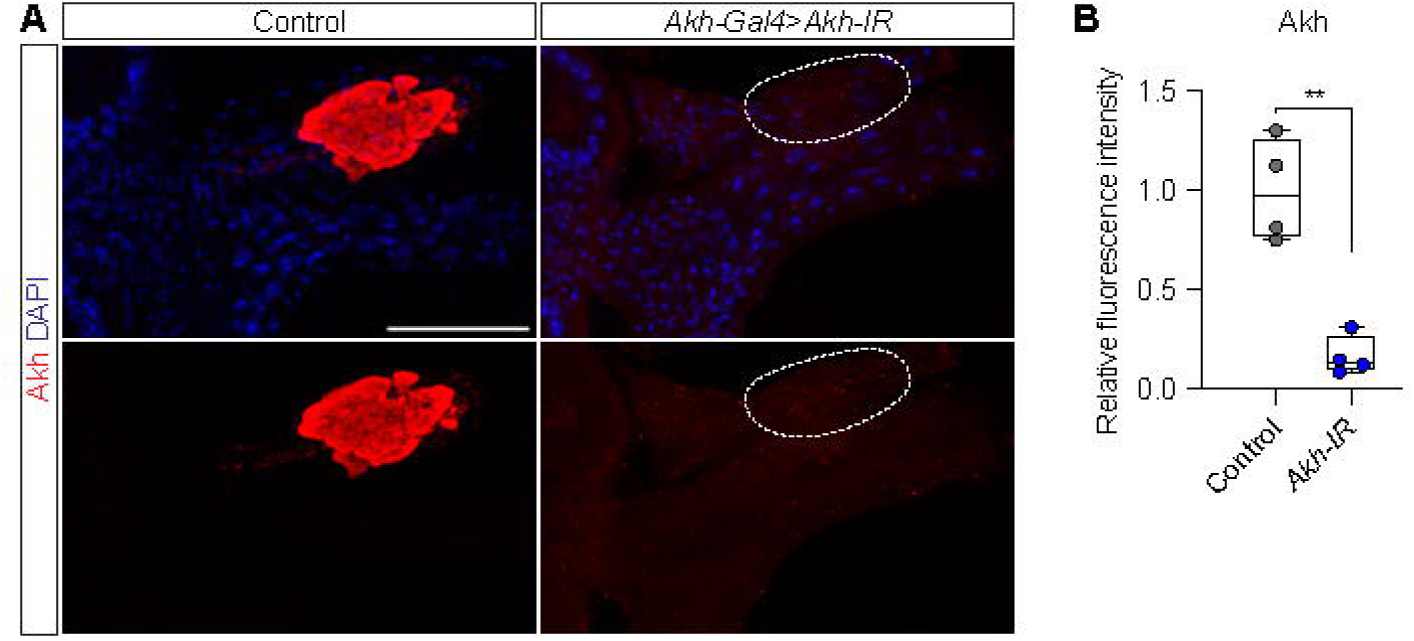
*Akh*-Gal4>*Akh*-IR depleted Akh. **(A)** Representative confocal images of APCs from female adults. Control, *Akh*-Gal4>*w*^1118^. Anti-Akh is in red. DAPI stains DNA in blue. Scale bar: 50 μm. **(B)** Quantitation of the relative fluorescence intensity of anti-Akh in A. n=4. Statistical analysis was performed using *t*-test corrected with Welch; **, P<0.01.

**Figure 5—figure supplement 1.**
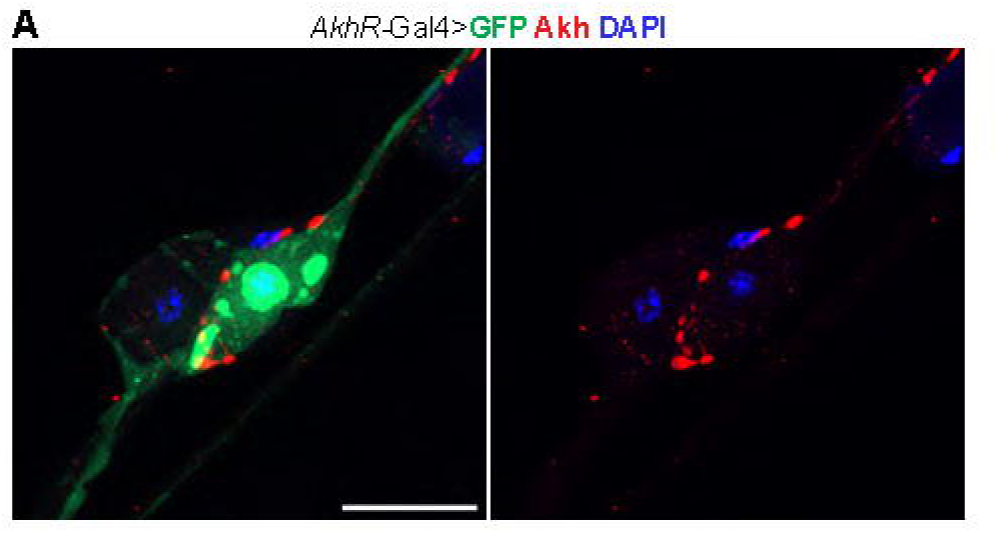
AkhR neuron receives Akh. (A) Representative confocal images of AkhR neuron from female adults (*AkhR*-Gal4>GFP). GFP is in green. Anti-Akh is in red. DAPI stains DNA in blue. Scale bar: 20 μm.

## REFERENCES

Agrawal S, Gupta Y. 2016. Comment on Kazda et al. Evaluation of Efficacy and Safety of the Glucagon Receptor Antagonist LY2409021 in Patients With Type 2 Diabetes: 12-and 24-Week Phase 2 Studies. Diabetes Care 2016;39:1241–1249. *Diabetes Care*.

Ambache N, Lippold OC. 1949. Bradycardia of central origin produced by injections of tetanus toxin into the vagus nerve. J Physiol 108:186–196.

Aranda-Domene R, Orenes-Piñero E, Arribas-Leal JM, Canovas-Lopez S, Hernández-Cascales J. 2023. Evidence for a lack of inotropic and chronotropic effects of glucagon and glucagon receptors in the human heart. Cardiovasc Diabetol 22:128.

Aune D, Feng T, Schlesinger S, Janszky I, Norat T, Riboli E. 2018. Diabetes mellitus, blood glucose and the risk of atrial fibrillation: A systematic review and meta-analysis of cohort studies. J Diabetes Complications 32:501–511.

Bharucha KN, Tarr P, Zipursky SL. 2008. A glucagon-like endocrine pathway in Drosophila modulates both lipid and carbohydrate homeostasis. J Exp Biol 211:3103–3110.

Birse RT, Choi J, Reardon K, Rodriguez J, Graham S, Diop S, Ocorr K, Bodmer R, Oldham S. 2010. High-fat-diet-induced obesity and heart dysfunction are regulated by the TOR pathway in Drosophila. Cell Metab 12:533–544.

Bomholt AB, Johansen CD, Christensen JB, Kjeldsen SAS, Galsgaard KD, Winther-Sørensen M, Serizawa R, Hornum M, Porrini E, Pedersen J, Ørskov C, Gluud LL, Sørensen CM, Holst JJ, Albrechtsen R, Wewer Albrechtsen NJ. 2022. Evaluation of commercially available glucagon receptor antibodies and glucagon receptor expression. Commun Biol 5:1278.

Cai C, Wu N, Yang G, Yang S, Liu W, Chen M, Po SS, TREAT PVC investigators. 2023. Transcutaneous electrical vagus nerve stimulation to suppress premature ventricular complexes (TREAT PVC): study protocol for a multi-center, double-blind, randomized controlled trial. Trials 24:683.

Chang-Chretien K, Chew JT, Judge DP. 2004. Reversible dilated cardiomyopathy associated with glucagonoma. Heart 90:e44.

D’Alessio D. 2011. The role of dysregulated glucagon secretion in type 2 diabetes. Diabetes Obes Metab 13 Suppl 1:126–132.

Ding M, Li QF, Yin G, Liu JL, Jan XY, Huang T, Li AC, Zheng L. 2022. Effects of Drosophila melanogaster regular exercise and apolipoprotein B knockdown on abnormal heart rhythm induced by a high-fat diet. PLoS One 17:e0262471.

Dulcis D, Levine RB. 2005. Glutamatergic Innervation of the Heart Initiates Retrograde Contractions in Adult Drosophila melanogaster. J Neurosci 25:271–280.

Dulcis D, Levine RB. 2003. Innervation of the heart of the adult fruit fly, Drosophila melanogaster. J Comp Neurol 465:560–578.

Dulcis D, Levine RB, Ewer J. 2005. Role of the neuropeptide CCAP in Drosophila cardiac function. J Neurobiol 64:259–274.

Farah A, Tuttle R. 1960. Studies on the pharmacology of glucagon. J Pharmacol Exp Ther 129:49–55.

Freeman AG. 1951. Electrocardiographic findings during operative manipulation of the viscera and vagus nerves. Lancet 1:926–930.

González ML, Pividori SM, Fosser G, Pontecorvo AA, Franco-Riveros VB, Tubbs RS, Boezaart AP, Reina MA, Buchholz B. 2023. Innervation of the heart: Anatomical study with application to better understanding pathologies of the cardiac autonomics. Clin Anat 36:550–562.

Grönke S, Müller G, Hirsch J, Fellert S, Andreou A, Haase T, Jäckle H, Kühnlein RP. 2007. Dual lipolytic control of body fat storage and mobilization in Drosophila. PLoS Biol 5:e137.

Gupta V, Munjal JS, Jhajj P, Jhajj S, Jain R. 2022. Obesity and Atrial Fibrillation: A Narrative Review. Cureus 14:e31205.

Habegger KM, Heppner KM, Geary N, Bartness TJ, DiMarchi R, Tschöp MH. 2010. The metabolic actions of glucagon revisited. Nat Rev Endocrinol 6:689–697.

Heppner KM, Habegger KM, Day J, Pfluger PT, Perez-Tilve D, Ward B, Gelfanov V, Woods SC, DiMarchi R, Tschöp M. 2010. Glucagon regulation of energy metabolism. Physiol Behav 100:545–548.

Huang R, Song T, Su H, Lai Z, Qin W, Tian Y, Dong X, Wang L. 2020. High-fat diet enhances starvation-induced hyperactivity via sensitizing hunger-sensing neurons in Drosophila. Elife 9:e53103.

Hurwitz RA. 1973. Effect of glucagon on infants and children with atrioventricular heart block. Br Heart J 35:1260–1264.

Huxley RR, Filion KB, Konety S, Alonso A. 2011. Meta-analysis of cohort and case-control studies of type 2 diabetes mellitus and risk of atrial fibrillation. Am J Cardiol 108:56–62.

Isabel G, Martin JR, Chidami S, Veenstra JA, Rosay P. 2005. AKH-producing neuroendocrine cell ablation decreases trehalose and induces behavioral changes in Drosophila. Am J Physiol Regul Integr Comp Physiol 288:R531–8.

Jaca IJ, Desai D, Barkin JS. 2002. Paroxysmal supraventricular tachycardia after administration of glucagon during upper endoscopy. Gastrointest Endosc 56:304.

Kannel WB, Abbott RD, Savage DD, McNamara PM. 1983. Coronary heart disease and atrial fibrillation: the Framingham Study. Am Heart J 106:389–396.

Kannel WB, Wolf PA, Benjamin EJ, Levy D. 1998. Prevalence, incidence, prognosis, and predisposing conditions for atrial fibrillation: population-based estimates. Am J Cardiol 82:2N–9N.

Kazda CM, Ding Y, Kelly RP, Garhyan P, Shi C, Lim CN, Fu H, Watson DE, Lewin AJ, Landschulz WH, Deeg MA, Moller DE, Hardy TA. 2016a. Evaluation of Efficacy and Safety of the Glucagon Receptor Antagonist LY2409021 in Patients With Type 2 Diabetes: 12- and 24-Week Phase 2 Studies. Diabetes Care 39:1241–1249.

Kazda CM, Ding Y, Kelly RP, Garhyan P, Shi C, Lim CN, Fu H, Watson DE, Lewin AJ, Landschulz WH, Deeg MA, Moller DE, Hardy TA. 2016b. Response to Comment on Kazda et al. Evaluation of Efficacy and Safety of the Glucagon Receptor Antagonist LY2409021 in Patients With Type 2 Diabetes: 12- and 24-Week Phase 2 Studies. Diabetes Care 2016;39:1241–1249. *Diabetes Care*.

Kharbanda RK, Ramdat Misier NL, van Schie MS, Zwijnenburg RD, Amesz JH, Knops P, Bogers AJJC, Taverne YJHJ, de Groot NMS. 2023. Insights Into the Effects of Low-Level Vagus Nerve Stimulation on Atrial Electrophysiology: Towards Patient-Tailored Cardiac Neuromodulation. JACC Clin Electrophysiol 9:1843–1853.

Kim SK, Rulifson EJ. 2004. Conserved mechanisms of glucose sensing and regulation by Drosophila corpora cardiaca cells. Nature 431:316–320.

Kodde IF, van der Stok J, Smolenski RT, de Jong JW. 2007. Metabolic and genetic regulation of cardiac energy substrate preference. Comp Biochem Physiol A Mol Integr Physiol 146:26–39.

Kusumoto FM, Schoenfeld MH, Barrett C, Edgerton JR, Ellenbogen KA, Gold MR, Goldschlager NF, Hamilton RM, Joglar JA, Kim RJ, Lee R, Marine JE, McLeod CJ, Oken KR, Patton KK, Pellegrini CN, Selzman KA, Thompson A, Varosy PD. 2019. 2018 ACC/AHA/HRS Guideline on the Evaluation and Management of Patients With Bradycardia and Cardiac Conduction Delay: Executive Summary: A Report of the American College of Cardiology/American Heart Association Task Force on Clinical Practice Guidelines, and the Heart Rhythm Society. Circulation 140:e333–e381.

Lee G, Park JH. 2004. Hemolymph sugar homeostasis and starvation-induced hyperactivity affected by genetic manipulations of the adipokinetic hormone-encoding gene in Drosophila melanogaster. Genetics 167:311–323.

Lee H, Choi E-K, Lee S-H, Han K-D, Rhee T-M, Park C-S, Lee S-R, Choe W-S, Lim W-H, Kang S-H, Cha M-J, Oh S. 2017. Atrial fibrillation risk in metabolically healthy obesity: A nationwide population-based study. Int J Cardiol 240:221–227.

Lee P-T, Zirin J, Kanca O, Lin W-W, Schulze KL, Li-Kroeger D, Tao R, Devereaux C, Hu Y, Chung V, Fang Y, He Y, Pan H, Ge M, Zuo Z, Housden BE, Mohr SE, Yamamoto S, Levis RW, Spradling AC, Perrimon N, Bellen HJ. 2018. A gene-specific T2A-GAL4 library for Drosophila. Elife 7. doi:10.7554/eLife.35574

Liao S, Amcoff M, Nassel DR. 2020. Impact of high-fat diet on lifespan, metabolism, fecundity and behavioral senescence in Drosophila. Insect Biochem Mol Biol 2020/11/11:103495.

Lucchesi BR, Brown J, Laidlaw B, Rockiki D. 1968. Cardiac Actions of Glucagon. Circ Res 22:777–787.

Markiewicz K, Cholewa M, Górski L. 1978. Cardiac arrhythmias after intravenous administration of glucagon. Eur J Cardiol 6:449–458.

Masuyama K, Zhang Y, Rao Y, Wang JW. 2012. Mapping neural circuits with activity-dependent nuclear import of a transcription factor. J Neurogenet 26:89–102.

Mukharji A, Drucker DJ, Charron MJ, Swoap SJ. 2013. Oxyntomodulin increases intrinsic heart rate through the glucagon receptor. Physiol Rep 1:e00112.

Nattel S, Guasch E, Savelieva I, Cosio FG, Valverde I, Halperin JL, Conroy JM, Al-Khatib SM, Hess PL, Kirchhof P, De Bono J, Lip GYH, Banerjee A, Ruskin J, Blendea D, Camm AJ. 2014. Early management of atrial fibrillation to prevent cardiovascular complications. Eur Heart J 35:1448–1456.

Neumann J, Hofmann B, Dhein S, Gergs U. 2023. Glucagon and Its Receptors in the Mammalian Heart. Int J Mol Sci 24. doi:10.3390/ijms241612829

Noyes BE, Katz FN, Schaffer MH. 1995. Identification and expression of the Drosophila adipokinetic hormone gene. Mol Cell Endocrinol 109:133–141.

Ocorr K, Reeves NL, Wessells RJ, Fink M, Chen H-SV, Akasaka T, Yasuda S, Metzger JM, Giles W, Posakony JW, Bodmer R. 2007. KCNQ potassium channel mutations cause cardiac arrhythmias in Drosophila that mimic the effects of aging. Proc Natl Acad Sci U S A 104:3943–3948.

Pearson MJ, Unger RH, Holland WL. 2016. Clinical Trials, Triumphs, and Tribulations of Glucagon Receptor Antagonists. Diabetes Care.

Powell-Wiley TM, Poirier P, Burke LE, Després J-P, Gordon-Larsen P, Lavie CJ, Lear SA, Ndumele CE, Neeland IJ, Sanders P, St-Onge M-P, Null N. 2021. Obesity and Cardiovascular Disease: A Scientific Statement From the American Heart Association. Circulation 143:e984–e1010.

Radulescu A, Gannon MC, Nuttall FQ. 2010. The effect on glucagon, glucagon-like peptide-1, total and acyl-ghrelin of dietary fats ingested with and without potato. J Clin Endocrinol Metab 95:3385–3391.

Roberts-Thomson KC, Lau DH, Sanders P. 2011. The diagnosis and management of ventricular arrhythmias. Nat Rev Cardiol 8:311–321.

Schneider CA, Rasband WS, Eliceiri KW. 2012. NIH Image to ImageJ: 25 years of image analysis. Nat Methods 9:671–675.

Scott RS, Espiner EA, Donald RA, Ellis MJ. 1980. Free insulin, C-peptide and glucagon profiles in insulin dependent diabetes mellitus. Aust N Z J Med 10:146–150.

Sowden GL, Drucker DJ, Weinshenker D, Swoap SJ. 2007. Oxyntomodulin increases intrinsic heart rate in mice independent of the glucagon-like peptide-1 receptor. Am J Physiol Regul Integr Comp Physiol 292:R962–70.

Staubli F, Jorgensen TJD, Cazzamali G, Williamson M, Lenz C, Sondergaard L, Roepstorff P, Grimmelikhuijzen CJP. 2002. Molecular identification of the insect adipokinetic hormone receptors. Proc Natl Acad Sci U S A 99:3446–3451.

Stern JH, Smith GI, Chen S, Unger RH, Klein S, Scherer PE. 2019. Obesity dysregulates fasting-induced changes in glucagon secretion. J Endocrinol 243:149–160.

Unger RH, Aguilar-Parada E, Müller WA, Eisentraut AM. 1970. Studies of pancreatic alpha cell function in normal and diabetic subjects. J Clin Invest 49:837–848.

van Weperen VYH, Vaseghi M. 2023. Cardiac vagal afferent neurotransmission in health and disease: review and knowledge gaps. Front Neurosci 17:1192188.

Vuguin PM, Charron MJ. 2011. Novel insight into glucagon receptor action: lessons from knockout and transgenic mouse models. Diabetes Obes Metab 13 Suppl 1:144–150.

Wang TJ, Parise H, Levy D, D’Agostino RB, Wolf PA, Vasan RS, Benjamin EJ. 2004. Obesity and the Risk of New-Onset Atrial Fibrillation. JAMA 292:2471–2477.

Yelbuz TM, Choma MA, Thrane L, Kirby ML, Izatt JA. 2002. Optical coherence tomography: a new high-resolution imaging technology to study cardiac development in chick embryos. Circulation 106:2771–2774.

Yu Y, Huang R, Ye J, Zhang V, Wu C, Cheng G, Jia J, Wang L. 2016. Regulation of starvation-induced hyperactivity by insulin and glucagon signaling in adult Drosophila. Elife 5:e15693.

Zhang K, Lehner LJ, Praeger D, Baumann G, Knebel F, Quinkler M, Roepke TK. 2014. Glucagonoma-induced acute heart failure. Endocrinol Diabetes Metab Case Rep 2014:140061.

Zhao Y, Johansson E, Duan J, Han Z, Alenius M. 2023. Fat- and sugar-induced signals regulate sweet and fat taste perception in Drosophila. Cell Rep 42:113387.

Zhao Y, Khallaf MA, Johansson E, Dzaki N, Bhat S, Alfredsson J, Duan J, Hansson BS, Knaden M, Alenius M. 2022. Hedgehog-mediated gut-taste neuron axis controls sweet perception in Drosophila. Nat Commun 13:7810.

